# The worm affair: Genetic diversity in two species of symbionts that co-occur in tubeworms from the Mediterranean Sea

**DOI:** 10.1101/2021.01.27.428081

**Authors:** Tal Zvi-Kedem, Eli Shemesh, Dan Tchernov, Maxim Rubin-Blum

## Abstract

The symbioses between the vestimentiferan tubeworms and their chemosynthetic partners (Gammaproteobacteria, Chromatiales, *Sedimenticolaceae*) hallmark the success of these organisms in hydrothermal vent and hydrocarbon seep deep-sea habitats. The fidelity of these associations varies, as both the hosts and the symbionts can be loose in partner choice. Some tubeworms may host distinct symbiont phylotypes, which often co-occur in a single host individual. To better understand the genetic basis for the ‘promiscuity’ of tubeworm symbioses, we curated and investigated metagenome-assembled genomes of two symbiont phylotypes (species, based on the average nucleotide identity <95%) in *Lamellibrachia anaximandri*, a vestimentiferan endemic to the Mediterranean Sea, in individuals collected from Palinuro hydrothermal vents (Italy) and hydrocarbon seeps (Eratosthenes seamount and Palmahim disturbance). Using comparative genomics, we show that mainly mobilome and defense mechanism-related features distinguish the symbiont genotypes. While many central metabolic functions are conserved in the tubeworm symbionts, nitrate respiration (Nar, Nap and Nas proteins) is modular, yet this modularity is not linked to speciation, but rather to local adaptation. Our results hint that variation in a single moonlighting protein may be responsible for the host-symbiont fidelity.

## Introduction

Siboglinid tubeworms (Polychaeta; Siboglinidae) are key taxa in deep-sea hydrothermal vents and hydrocarbon seeps (McMullin *et al.*, 2003; Bright *et al.*, 2013). These animals owe their success in these chemosynthetic habitats to symbioses with sulfide-oxidizing gammaproteobacteria (Bright and Lallier, 2010). These bacteria are classified as members of the *Sedimenticolaceae* family (Chroamtiales) by the Genome Taxonomy Database (Parks *et al.*, 2018; Rinke *et al.*, 2020), thus share a recent common ancestor with the photoautotrophic purple sulfur bacteria. There is genomic, transcriptomic and proteomic evidence for adaptation of these highly efficient autotrophs to the symbiotic lifestyle in a chemosynthetic environment (Markert *et al.*, 2007, 2011; Gardebrecht *et al.*, 2012; Li *et al.*, 2018; Hinzke, Kleiner, Breusing, Felbeck, Häsler, Stefan M Sievert, *et al.*, 2019; M. Rubin-Blum *et al.*, 2019; Yang *et al.*, 2019; Hinzke *et al.*, 2021).

Vestimentiferan tubeworms acquire their horizontally-transmitted, intracellular symbionts from a genetically-diverse pool of free-living lineages in the surrounding environment each generation (Nussbaumer *et al.*, 2006; Klose *et al.*, 2015; Polzin *et al.*, 2019). The fidelity of these associations is often high, as usually a single host species, e.g. *Riftia pachyptila*, carry a single symbiont phylotype, in this case, *Candidatus* Endoriftia persephone (Bright *et al.*, 2013). However, these symbionts are ‘promiscuous’ with regards to a host, as *Ca.* Endoriftia persephone also occupies the trophosomes of *Tevnia jerichonana* and *Ridgeia piscesae.* This is also the case for the symbiont phylotypes that are hosted by the seep tubeworms *Lamelibrachia luymesi* and *Seepiophila jonesi* (Li *et al.*, 2018). It is now evident that symbiont species comprise multiple strains, although frequency-dependent selection or preferential growth following unspecific uptake may limit the intra-species strain-level symbiont diversity (Bright and Bulgheresi, 2010; Vrijenhoek, 2010; Perez and Juniper, 2016; Polzin *et al.*, 2019; Russell, 2019). Bottlenecks during symbiont transmission and host colonization can greatly reduce the diversity of symbionts within each host individual (Bright and Bulgheresi, 2010; Vrijenhoek, 2010; Russell, 2019).

Yet, the hosts can also be ‘promiscuous’ and establish partnerships with more than one symbiont lineage. Two distinct symbiont phylotypes were found in tubeworms from the *Lamellibrachia* genus: amplicon and Sanger sequencing, as well as fluorescence in situ hybridization (FISH), revealed that *L. barhami* from the cold seeps in the Eastern Pacific, as well as *L. anaximandri*, which is endemic to the Mediterranean Sea (Southward *et al.*, 2011), often host more than one symbiont phylotype, based on the dissimilarities in the 16S rRNA genes of the symbionts (Rubin-Blum *et al.*, 2014; Zimmermann *et al.*, 2014; Breusing, Franke, *et al.*, 2020). On one hand, this is unusual, especially since the competition among symbionts may be unfavorable for the fitness of the host (Frank, 1996). On the other hand, the economy of the low-cost chemosynthetic symbioses, in which the environment provides nutrition, differs from that of symbioses in which the host is obliged to ‘feed’ the symbionts, thus the chemosynthetic symbioses have a higher tolerance for symbiont diversity (Ansorge *et al.*, 2019). For example, multiple strains of often functionally-diverse sulfur-oxidizing symbionts co-exist in bathymodiolin mussels (Ikuta *et al.*, 2015; Ansorge *et al.*, 2019).

Here we asked if there are potential functional differences in distinct symbiont phylotypes that occupy the same host species, and how the genomic variability could influence the fitness of these symbioses. For that, we sequenced and assembled high-quality genomes of the two *L. anaximandri* symbiont phylotypes and explored their phylogeny and pangenomes. It has been suggested that the geographic structuring of the free-living symbiont population plays a major role in forming the microbiome in some deep-sea chemosynthetic associations, such as those of bathymodiolin mussels (Ücker *et al.*, 2020). We thus further investigated genomic variation among symbiont species found in different geographic locations and habitats - Palinuro hydrothermal vents in the Tyrrhenian Sea, as well as hydrocarbon seeps at the Eratosthenes seamount and Palmahim disturbance in the eastern Mediterranean Sea (**Fig. 1**), following a previous diversity study (Rubin-Blum *et al.*, 2014). We also hypothesized that in the very warm (~15°C) deep waters of the Mediterranean Sea, the symbionts may have acquired distinct traits, and thus compared the genomes of *L. anaximandri* symbionts to those of other hosts.

**Figure 1:**
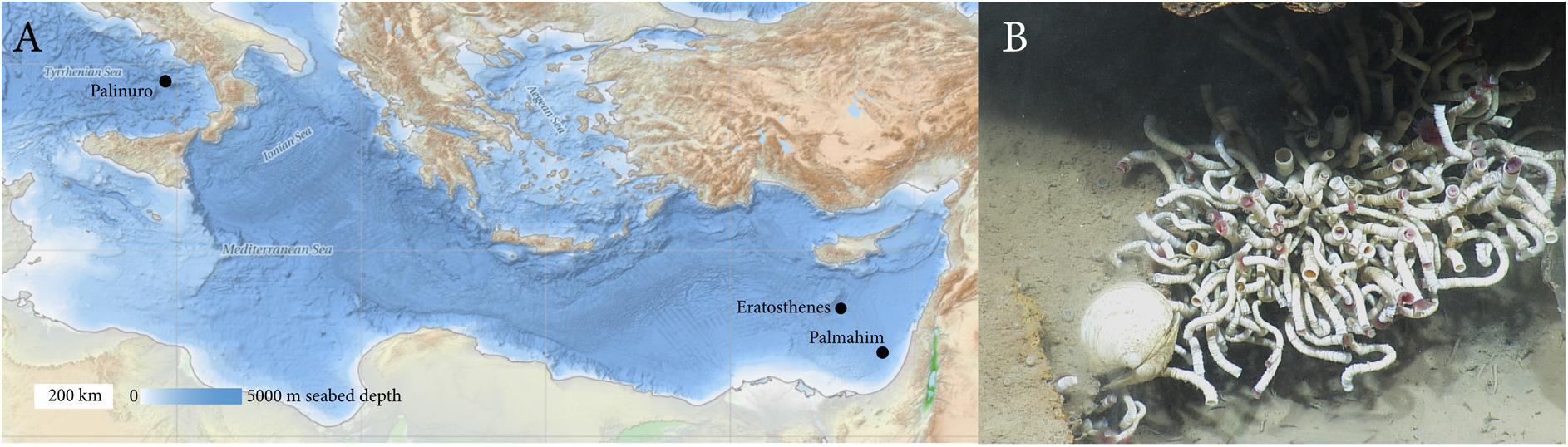
*Lamellibrachia anaximandri* samples collected in this study. A) Map showing the collection sites. B) *L. anaximandri* colony at the Palmahim disturbance. The bathymetric map was downloaded from the EMODnet Bathymetry portal - http://www.emodnet-bathymetry.eu. The image was acquired by the during the NA19 Nautilus *E/V* expedition and is a courtesy of the Ocean Exploration Trust.

## Results and Discussion

### *L. anaximandri* hosts two distinct symbiont species

We curated five high-quality metagenome-assembled genomes (MAGs, 99% completeness and 1-2% contamination, **Supporting Information Table 1**), from a total of 9 metagenomic libraries (two distinct phylotypes from Palmahim individuals, a single phylotype from Eratosthenes individuals, and a single phylotype from Palinuro - two MAGs from distinct individuals to account for intra-species variation). Sample collection metadata has been described in detail previously (Rubin-Blum *et al.*, 2014), and summarized in **Fig. 1**. Although two distinct phylotypes co-occurred in a Palamhim individual, as detected by phylotype-specific PCR (Rubin-Blum *et al.*, 2014) and binning, as shown in **Supplementary Information Fig. S1**, we were unable to curate more than one MAG from the same library, due to insufficient coverage of the low-abundance phylotype. Phylotype A MAGs (16S rRNA sequence identical to that of type A clone, GeneBank accession KC832733.1, Zimmerman *et al.* 2014) were binned from Palinuro and Palmahim individuals, while those of phylotype B (99.6% identity of the 16S rRNA sequence to that of type B clone, GeneBank accession KC832729.1, Zimmerman *et al.* 2014) were binned from Palmahim and Eratosthenes individuals.

We defined phylotypes A and B as distinct species. This is based on the 93% average nucleotide identity (ANI) for all the phylotype A and B genome pairs (**Fig. 2**) and the 95% ANI boundary for bacterial species (Jain *et al.*, 2018). Phylogenomic analysis of 159 conserved proteins suggests that *L. anaximandri* symbionts fall into the ‘seep’ sister clade of *Ca.* Endoriftia persephone (**Fig. 2**). We note that *Ca.* Endoriftia persephone and bacteria from the ‘seep’ clade appear to comprise the same genus, based on the 61 ±3 percentage of conserved proteins (POCP), which is larger than the minimum 50% POCP boundary for all the lineages within a genus (Qin *et al.*, 2014). While species B genotypes from Eratosthenes and Palmahim were highly similar (99.9%ANI), species A genotypes differed between Palinuro and Palmahim individuals, as suggested by phylogeny, and 98.1% ANI (**Fig. 2**). This hints at the geographic structuring of the symbiont populations, yet a larger-scale study is needed to further test this hypothesis. We found no clear linkage between the host mitochondrial genotype and the presence of specific symbiont lineages (**Supplementary Information Fig. S2**). Our results thus suggest that no symbiont selection mechanism is linked to the mitochondrial genotype in *L. anaximandri*, however, this doesn’t rule out the possibility that traits encoded by the nuclear genome of the host may select for one or the other symbiont.

**Figure 2.**
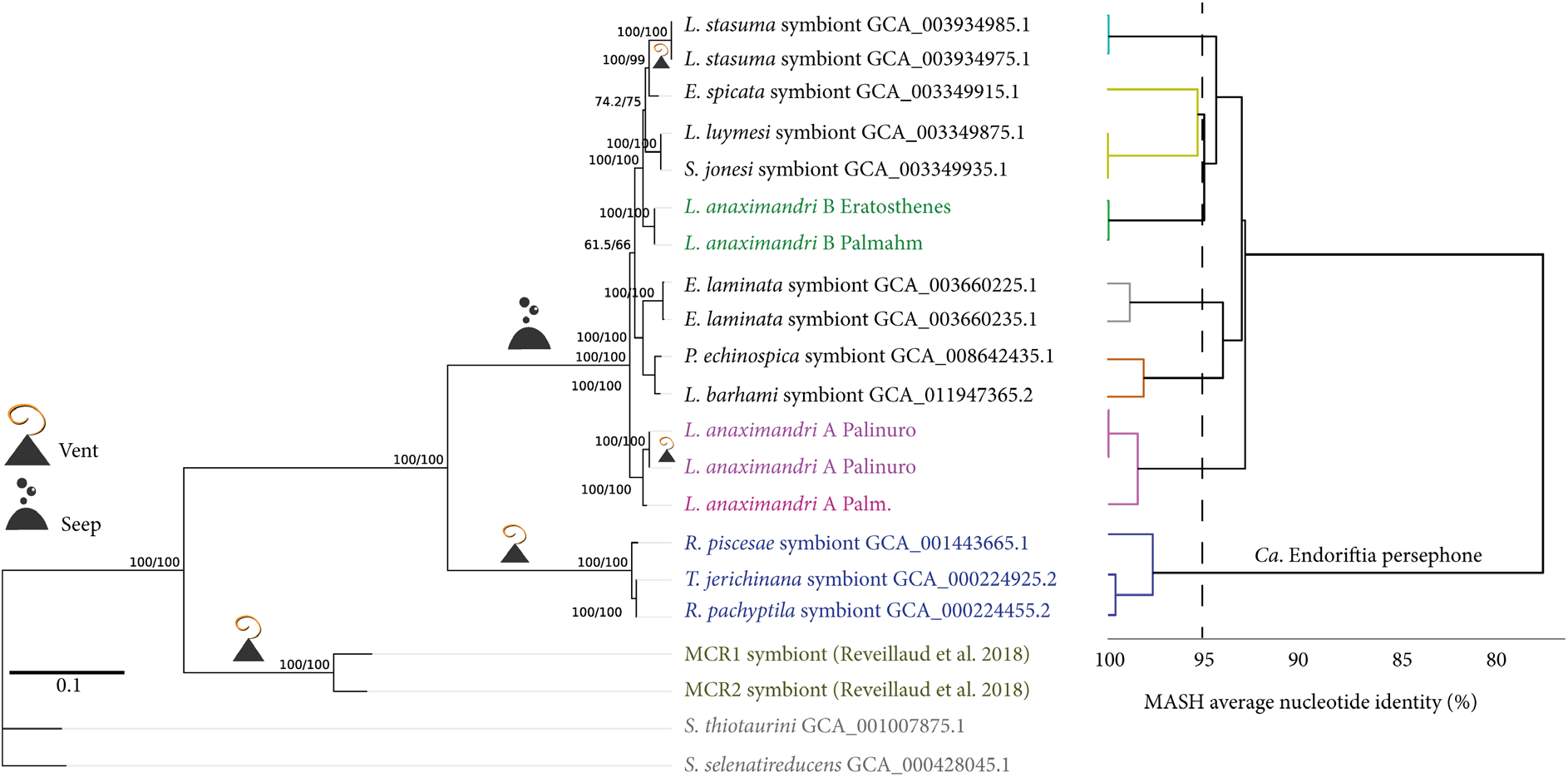
Phylogenomic tree of vestimentiferan tubeworm symbionts and the related free-living lineages, based on the alignment of 159 conserved proteins. Average nucleotide identity values and their clustering are shown, indicating the 95% cutoff between species (different colors). Shimodaira–Hasegawa approximate likelihood-ratio test (SH-aLRT) and ultrafast bootstrap approximation (UFBoot) branch support values are shown next to the branches. Average nucleotide identity (ANI) values determined by Mash and their clustering are shown on the right, and colors indicate distinct species based on the 95% species boundary (See Supplementary Information Fig. S5 for Anvi’o representation of this clade.).

**Figure 3:**
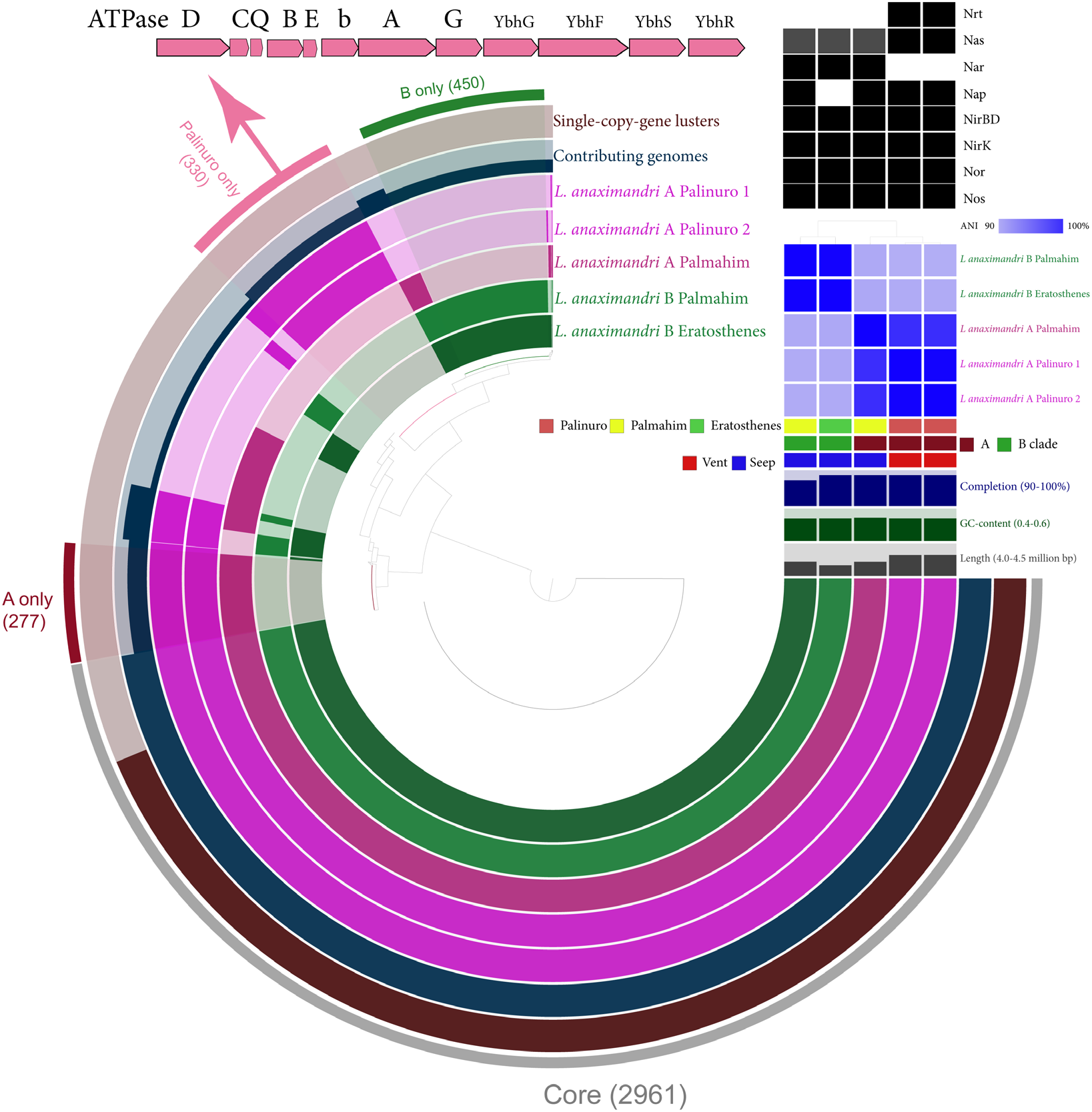
The comparison of five *Lamellibrachia anaximandri* symbiont genomes. In the circular representation, orthologous gene clusters are clustered by their absence/presence in the genomes. Genome statistics (completeness, GC content and length), metadata, ANI heatmap and clustering are shown. A grayscale heatmap in the right corner describes the absence (white) and presence (grey or black, for distinct alleles) of features in nitrate reduction steps. We also show the gene cluster that encodes the DCQBEbAG subunits of a sodiumtransporting ATPase, together with an ATP-dependent ABC transporter YbhGFSR, found only in Palinuro symbionts.

### Biotic interactions and modularity of nitrate metabolism shape the accessory genomes

Comparisons of the *L. anaximandri* symbiont MAGs revealed 2961 core genes (**Fig. 2**). The core genomes encoded the enzymes needed to sustain the complex metabolism of the chemosynthetic tubeworm symbionts, which has been described in detail (Robidart *et al.*, 2008; Li *et al.*, 2018; Yang *et al.*, 2019; Breusing, Schultz, *et al.*, 2020). These encoded functions allow the symbionts to fix inorganic carbon via Calvin and reductive Krebs cycles, oxidize sulfide and store elemental sulfur, assimilate nitrate and ammonia, denitrify to produce N2 and potentially interact with the host. In most cases concerning the central metabolism, we observed functional homogeneity between the different species of *L. anaximadri* symbionts, as well as within the whole seep clade.

We, however, identified marked differences in the genome content of the distinct symbiont genotypes. 277 and 450 unique gene clusters were attributed to species A and B, respectively. We also found 330 unique gene clusters in the genomes of species A from Palinuro (these were absent from the species-A genome from Palmahim). Analysis of Clusters of Orthologous Genes (COG) category enrichment indicated that functions related to mobilome (X) and signal transduction mechanisms (T) (**Supplementary Information Fig. S3**) were characteristic of the accessory genomes. We note that signal transduction mechanisms are often found in phages (Hargreaves *et al.*, 2014) and usually comprise phage defense mechanisms, such as restriction-modification, phage growth limitation and abortive infection systems (Depardieu *et al.*, 2016; Hampton *et al.*, 2020). We also found unique gene clusters that were linked to either Eastern Mediterranean (Palmahim + Eratosthenes) or Tyrrhenian Sea (Palinuro) locations. Most notably, 450 gene clusters were found only in the species A MAGs from Palinuro. These 450 gene clusters were enriched in defense mechanisms (V), replication, recombination and repair (L) and transcription (K) functions (**Supplementary Information Fig. S3**), which are most often attributed to phage predation and defense. Overall, the accessory genomes carried features such as transposases, phage integrases, toxin-antitoxin systems as well as CRISPR-associated proteins and restriction-modification systems (See **Supplementary Information Table. S2** for the EGGnog 5 annotations). The fact that the accessory genomes were enriched in these features is not surprising in light of models that explain the preservation of prokaryotic diversity of prokaryotic by phage predation (Rodriguez-Valera *et al.*, 2009, 2016).

Following a growing understanding of nitrate respiration modularity in symbiotic and free-living bacteria (Kleiner *et al.*, 2012; Ansorge, Romano, Sayavedra, Porras, *et al.*, 2019), one of the main discrepancies between the symbiont genotypes was in gene clusters needed for nitrate respiration and assimilation (**Fig. 2**). Only the type A symbionts encoded the three subunits of the ABC nitrate transporter, which are not found in the other tubeworm symbionts. Most strikingly, only the Palmahim B genotypes lacked the *napH* and *napG* genes of the periplasmic, high-affinity nitrate reductase Nap, which is otherwise found in the genomes of all the vestimentiferan tubeworm symbionts explored here. Read mapping confirmed that this was not a binning artifact, and some remnants of the Nap gene cluster were found in Palmahim B MAGs (*napC* and partial *napB*). The respiratory nitrate reductase Nar, which sporadically occurs in tubeworm symbionts (Breusing, Schultz, *et al.*, 2020), was found only in the genomes Eratosthenes and Palmahim symbionts (both A and B, *narQKGHIJ).* Bacteria use the high-affinity Nap under nitrate limitation, while Nar, which is more favorable for ATP synthesis, is upregulated at high (mM) nitrate concentrations (Moreno-Vivián *et al.*, 1999; Kraft *et al.*, 2011). Low nitrate concentrations in hydrothermal fluids, such as those of Palinuro, may indeed select for the high-affinity nitrate reductases (Vetriani *et al.*, 2014). *Riftia pachyptila* can concentrate nitrate in blood, and excess nitrate occurs in it as a result of limited sulfide supply and thus reduced metabolism (Hahlbeck *et al.*, 2005). Thus, nitrate availability to the symbionts is likely determined by several parameters, including the ambient nitrate and sulfide concentrations, as well as the host physiology, which scale the merit of losing or maintaining high and low-affinity nitrate uptake and reduction mechanisms.

We found that some features differed between the genotypes not only at the level of gene cluster presence/absence but also sequence similarity. All the symbionts carry the assimilatory nitrate reductase Nas, most similar to the NasCB form in *Bacillus subtilis*, based on the presence of two iron-sulfur clusters in the small subunit (Moreno-Vivián and Flores, 2007). The sequences of the small subunit differed markedly between *L. anaximadri* symbionts from the eastern Mediterranean and those from Palinuro (~80% sequence identity) and the phylogeny of these genes reveals two distinct clusters of symbiont-related sequences (**Supplementary Information Fig. S4**). Together with the absence of Nar in Palinuro symbionts, this considerable variation at the protein level likely results in the plasticity of nitrate uptake.

Comparative genomics revealed two functional gene clusters that differed among the genotypes and are not involved directly in nitrogen metabolism. Only the Palinuro genotypes (species A) carried a circa 13000 bp long gene cluster that encoded the DCQBEbAG subunits of a sodium-transporting ATPase, together with an ATP-dependent ABC transporter YbhGFSR (**Fig. 2**). This feature was absent from the genomes of other symbiotic and free-living relatives investigated here. The presence of an adjacent phage integrase indicates the potential for acquisition of the ATPase and ABC transporter via horizontal gene transfer. In *Escherichia coli*, Ybh transporters excrete tetracyclines, but also confer tolerance to NaCl, LiCl, and an alkaline pH (Feng *et al.*, 2020), and thus may allow adaptation for specific physiological conditions. Only species B symbionts carried a gene cluster involved in fatty acid metabolism, including genes that encoded (in this order) very long-chain acyl-CoA dehydrogenase, 3-ketoacyl-CoA thiolase and 3-hydroxyacyl-CoA dehydrogenase (FadAB), as well as FixAB electron transfer flavoprotein, indicating the ability to use alternative carbon sources (Srivastava *et al.*, 2015; Menendez-Bravo *et al.*, 2017).

### Omp porins may be responsible for the specificity of symbiont-host interactions

We further asked if there are gene clusters that may be responsible for the fidelity of the symbionts to the *L. anaximandri* hosts. We identified 20 gene clusters that were found only in *L. anaximandri* symbionts, but not in other ‘seep’ clade lineages (40 clusters if considering 80% sensitivity, that is, the gene was absent in the negatives, but present only in 4 out of 5 positives, see **Supplementary Information Fig. S5** for the clade-level Anvi’o analyses). We found no gene clusters specific to all other ‘seep’ clade lineages, but absent in *L. anaximandri* symbionts. A large fraction of the ‘*L. anaximandri* symbiont’-specific genes were poorly defined, and some functions encoded, such as nitrilase and arylsulfatase, are difficult to interpret. Only in the *L. anaximandri* symbionts, we identified a second aberrant copy of the *soxYZ* gene tandem, which encodes the protein complex that carries the intermediates of the sulfur-oxidizing system Sox (Sauvé *et al.*, 2007). The traits that account for the adaptation of *L. anaximandri* symbioses to the unusually warm deep water of the eastern Mediterranean Sea (as well as that of Palinuro hydrothermal vents) remain to be determined.

Most interestingly, we found that OmpA proteins, which encode a porin, were conserved with 98% identity in *L. anaximandri* symbionts (both phylotypes), but differed markedly from those of the closest hit within the ‘seep’ clade (44%). These porins, which are known to ‘moonlight’ (Jeffery, 2003), were especially highly expressed by the symbiont populations of *R. pachyptila* at protein level (Hinzke *et al.*, 2019), and *P. echinospica* and *E. laminata* at the RNA level (Rubin-Blum *et al.*, 2019; Yang *et al.*, 2019), indicating their central role in the symbiont physiology. We thus provide an additional line of evidence that such porins can be involved in the host-symbiont interactions and possibly determine their fidelity.

### Conclusions

Our results suggest that different forces shape the genomes of tubeworm symbionts. A history of phage predation is largely accountable for the pangenome expansion. Some of these features appear to escape streamlining, being integrated into defense mechanisms (Stern and Sorek, 2011) or due to the presence of toxin-antitoxin systems (Van Melderen, 2010). Horizontal gene transfer (e.g. YbhGFSR-ATPase), environmental selection (e.g. Nap and Nar) and genome streamlining result in a mosaic of traits that distinguish the native genotypes. These traits are not necessarily linked to the long-term evolutionary history of the symbiont lineages, and not to speciation. In particular, symbiont speciation is not linked directly to the fidelity of the symbiont-host associations. ‘Moonlighting’ porins, which are strongly diverged between the symbionts from different hosts and often highly expressed, may alone be accountable for the symbiont-tubeworm recognition, yet this remains to be confirmed experimentally. Future studies are also needed to elucidate the full scale of genomic diversity of these symbioses in the Mediterranean Sea and worldwide.

## Experimental procedures

DNA was extracted using PowerSoil kit (Qiagen) from trophosome subsections of eight *L. anaximandri* individuals, collected from the Eratosthenes seamount (33°38’ N, 32°47’ E, 947 m water depth), Palinuro volcanic complex (39°32’ N, 14°42’ E, 618 m water depth) and at the toe of the Palmachim disturbance (32°10’ N and 34°10 ‘E, 1036 m water depth). The presence of symbiont phylotypes was confirmed with PCR as previously described (Rubin-Blum *et al.*, 2014), using the primer set 27f-CM: 5’-AGAGTTTGATCMTGGCTCAG-3, Phylotype A: 5’-TCCTGCATCTCTCTGCTGGATTCTGTCA-3’ and Phylotype B: 5’-CCTCAGAACTTGTTAGAGATAACTTG-3’. Library preparation and sequencing of 30-60 million 150×2 bp Illumina HiSeq reads per individual was performed at HyLabs, Israel.

### Bioinformatics

Metagenomes were assembled using SPAdes V3.14 (Prjibelski *et al.*, 2020), following read preparation with the BBtools suite (Bushnell, B, sourceforge.net/projects/bbmap/). BBtools suite was also used for mapping of reads against the genomes, to estimate the number of reads mapped to features. MAGs were binned based on coverage, GC content and taxonomy with gbtools (Seah and Gruber-Vodicka, 2015), scaffolds < 800 bp were removed and genome completeness was estimated with CheckM (Parks *et al.*, 2015).

Phylogenomics were performed on 9897 parsimony-informative sites using the LG+F+R2 model with IQ-TREE 2 (Minh *et al.*, 2020), based on a concatenation of 159 conserved amino acid sequences determined with GTOtree (Lee and Ponty, 2019), in which poorly aligned positions were eliminated with Gblocks (Talavera and Castresana, 2007). Host phylogeny was constructed based on the alignment of mitochondrial genomic DNA sequence (11580 bp out of circa 15000 complete mitochondrial sequences). Mitochondrial genomes were assembled with mitoZ (Meng *et al.*, 2019), rotated with MARS (Ayad and Pissis, 2017) and aligned with MAFFT (Katoh and Standley, 2013). The tree was constructed with IQ-TREE 2 using TN+F model. Poorly aligned edges were manually removed.

Genomes were compared and gene clusters were defined with Roary (Page *et al.*, 2015), using -i 70 -cd 95 parameters and with Anvi’o V6.2 (Eren *et al.*, 2015), using --mcl-inflation 6 parameter. We used Scoary to examine trait associations (Brynildsrud *et al.*, 2016). To ensure that binning had little effect on the accessory genome estimates, we counted the coverage of MAG features with featureCounts (Liao *et al.*, 2014) following read mapping with 0.8 identity threshold and normalized the counts to those of the *cbbM* gene, following another normalization step to gene length. COG categories were assigned and quantified via DIAMOND BLAST (Buchfink *et al.*, 2014) search against the NCBI’s COG2014 database (Tatusov *et al.*, 2000) and the DIAMOND_COG_analysis_counter.py script (https://github.com/transcript/COG). We used PAST V4.03 (Hammer *et al.*, 2001) for the principal component analysis. ANI was determined with PyANI (Pritchard *et al.*, 2016) and Mash (Ondov *et al*, 2016). POCP was calculated with CompareM (https://github.com/dparks1134/CompareM) as described previously (Orata *et al.*, 2018).

## Data availability

Symbiont metagenome-assembled genomes were deposited to NCBI with project accession number PRJNA692100.

## Author contribution

T.Z-K, M.R-B and D.T conceived this study. D.T. and M.R-B organized and conducted sample collection. M.R-B, E.S. and T. Z-K performed DNA extractions and PCR. M.R-B was responsible for the bioinformatics. M. R-B wrote the paper with contributions from all co-authors.

## Acknowledgments

This research used samples and data provided by the E/V Nautilus Exploration Program: expeditions NA008, NA009, NA015 and NA019. The authors would like to thank all individuals who helped during the expedition, including onboard technical and scientific personnel, and the captain and crew of the E/V Nautilus. This study is funded by the Israeli Science Foundation (ISF) grant 913/19 to MRB and the Morris Kahn Marine Research Station.

## Supplement

**Table S1:** Genomes and metagenome-assembled genomes (MAGs) used in this study. Completeness (Comp.), contamination (Cont.), strain heterogeneity (SH) as determined by CheckM, genome size and N50 (from Quast) are shown.

| Symbiont MAG<br>(Host, genotype) | GeneBank<br>assembly accession | Comp.<br>(%) | Cont.<br>(%) | SH<br>(%) | Size<br>10<sup>6</sup> bp | N50<br>10<sup>4</sup> bp |
| --- | --- | --- | --- | --- | --- | --- |
| *L. anaximandri* A Palmahim | JAESPB000000000 | 98.92 | 2.04 | 0 | 4.22 | 4.66 |
| *L. anaximandri* A Palinuro 1 | JAESPC000000000 | 98.98 | 2.03 | 37.5 | 4.26 | 5.12 |
| *L. anaximandri* A Palinuro 2 | JAESPD000000000 | 98.98 | 1.22 | 0 | 4.26 | 5.34 |
| *L. anaximandri* B Eratosthenes | JAESPE000000000 | 99.33 | 2.7 | 0 | 4.14 | 3.25 |
| *L. anaximandri* B Palmahim | JAESPF000000000 | 99.33 | 2.7 | 0 | 4.20 | 3.30 |
| *E. laminata* A | GCA_003660225.1 | 99.32 | 1.52 | 0 | 4.20 | 10.61 |
| *E. laminata* B | GCA_003660235.1 | 99.06 | 1.4 | 0 | 4.25 | 12.50 |
| *E. spicata* | GCA_003349915.1 | 98.98 | 2.29 | 0 | 4.06 | 31.35 |
| *L. luymesii* | GCA_003349875.1 | 97.07 | 1.39 | 20 | 3.52 | 2.06 |
| *L. barhami* | GCA_011947365.2 | 96.57 | 2.46 | 80 | 4.17 | 414.8 |
| *R. piscesae* | GCA_001443665.1 | 98.83 | 0.7 | 0 | 3.44 | 9.89 |
| *R. pachyptila* | GCA_000224455.2 | 95.07 | 1.74 | 0 | 3.48 | 2.97 |
| *S. jonesi* | GCA_003349935.1 | 98.11 | 1.05 | 0 | 3.53 | 2.06 |
| *T. jerichonana* | GCA_000224925.2 | 98.35 | 1.05 | 0 | 3.64 | 9.27 |
| *P. echinospica* | GCA_008642435.1 | 97.24 | 2.64 | 46.15 | 4.06 | 38.16 |
| *L. satsuma* | GCA_003934985.1 | 98.95 | 2.73 | 0 | 3.91 | 7.30 |
| *L. satsuma* | GCA_003934975.1 | 97.79 | 2.73 | 0 | 3.93 | 6.94 |
| MCR1 | Reveillaud et al.<br>2018 | 97.95 | 2.22 | 0 | 4.26 | 8.99 |
| MCR2 | Reveillaud et al.<br>2018 | 98.52 | 4.11 | 16.67 | 4.64 | 4.27 |
| Free-living (genus, species) |  |  |  |  |  |  |
| *S. selenatireducens* | GCA_000428045.1 | 99.77 | 0.55 | 0 | 4.59 | 23.23 |
| *S. thiotaurini* | GCA_001007875.1 | 99.77 | 1.03 | 0 | 3.96 | 392.9 |

Table S2 (excel file): EGGnog annotations of proteins encoded by the core and accessory genomes: species A and B, Palinuro genotype.

**Figure S1:**
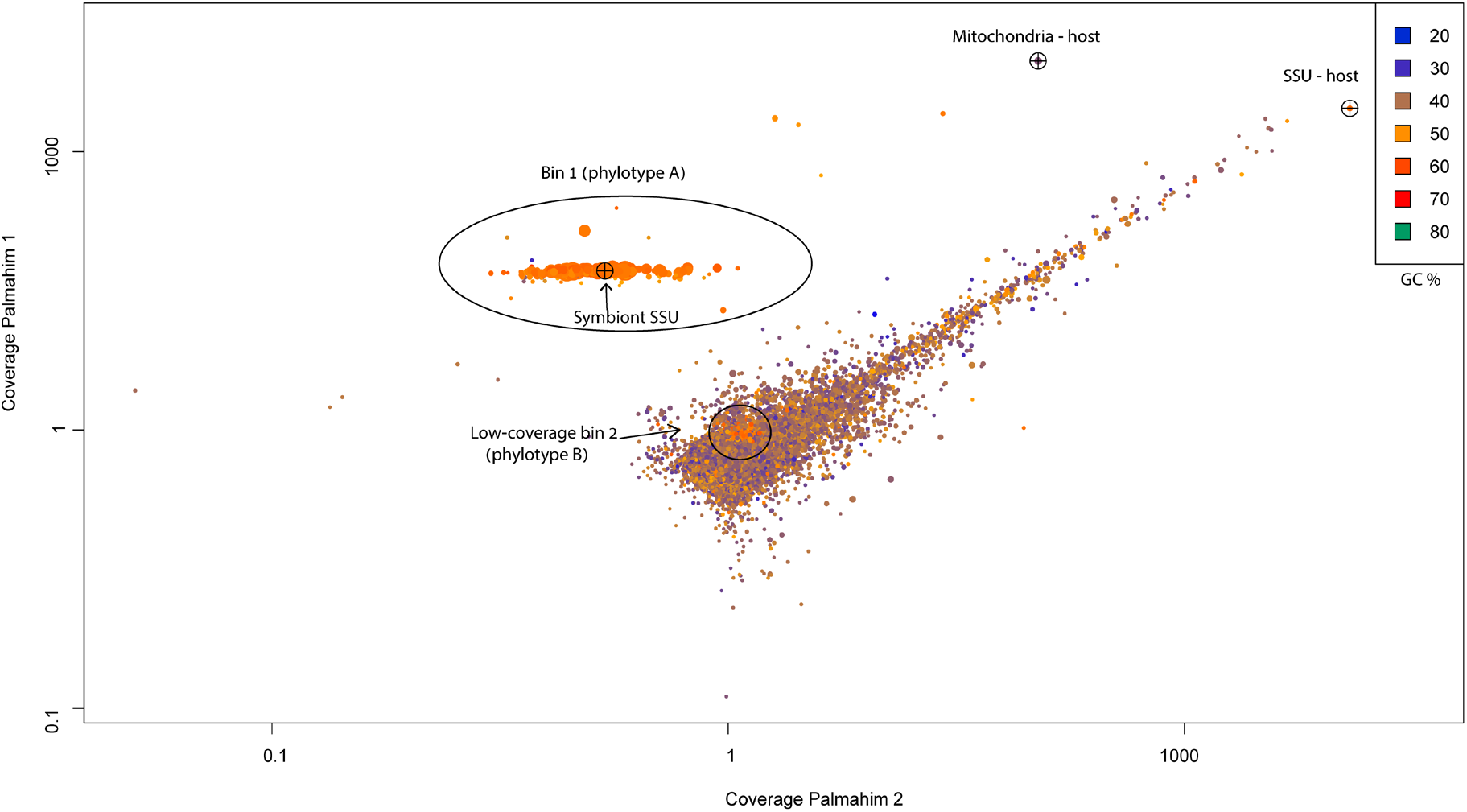
Differential coverage plot two metagenomic libraries from Palamhim. Scaffolds are represented by a point, whose diameter is proportional to the length of the scaffold. The main bin (phylotype A MAG) and the low coverage scaffolds from the co-occurring type B symbiont are shown. Points are colorized according to the GC content. The plot was produced by gbtools.

**Figure S2:**
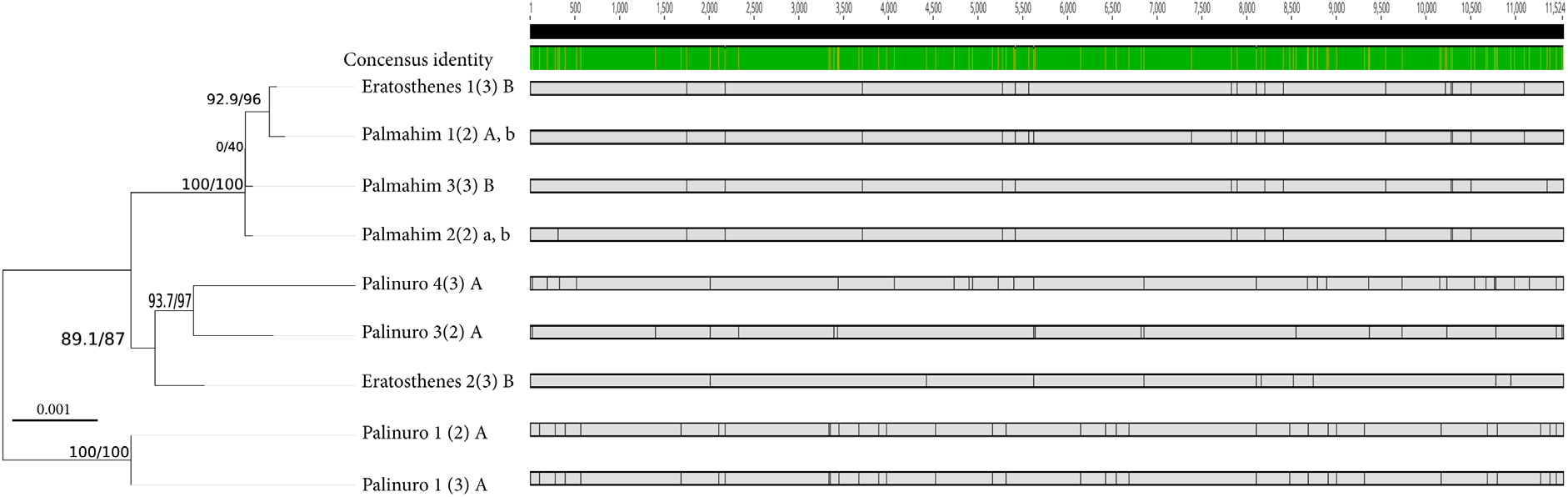
Phylogeny of the *L. anaximandri* mitochondrial sequences and single nucleotide polymorphisms. For each mitochondrial sequence, location, individual serial number and section number (in parentheses) are shown, as well as the symbiont phylotypes (A or B, capital letters represent the highly abundant phylotype). The fact that no mitochondrial genotype is specific to the geographic location indicates that there is no genetic barrier between the *L. anaximandri* populations.

**Figure S3:**
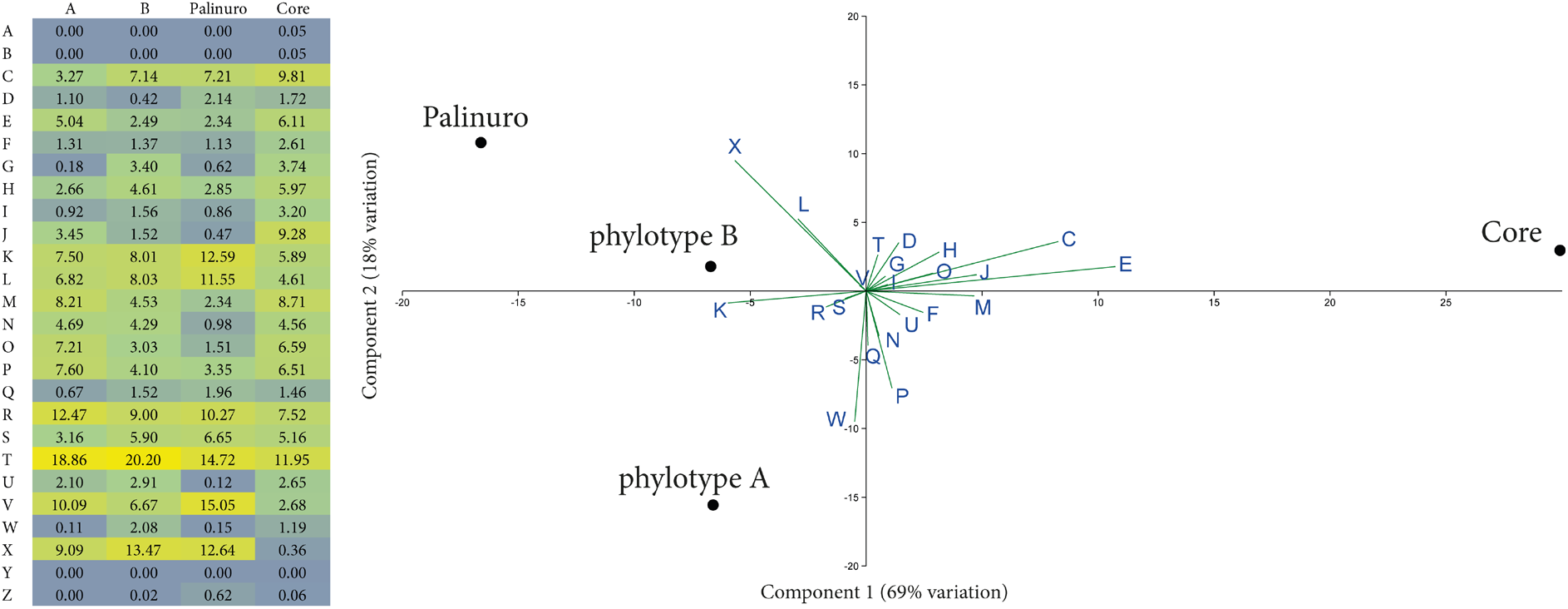
Clusters of orthologous groups (COG) functions attributed to the core genome, as well as to the accessory genomes of phylotypes A and B, as well as that of Palinuro genotype (phylotype A). Heatmap (left) and principal component analysis (right) are shown.

**Figure S4:**
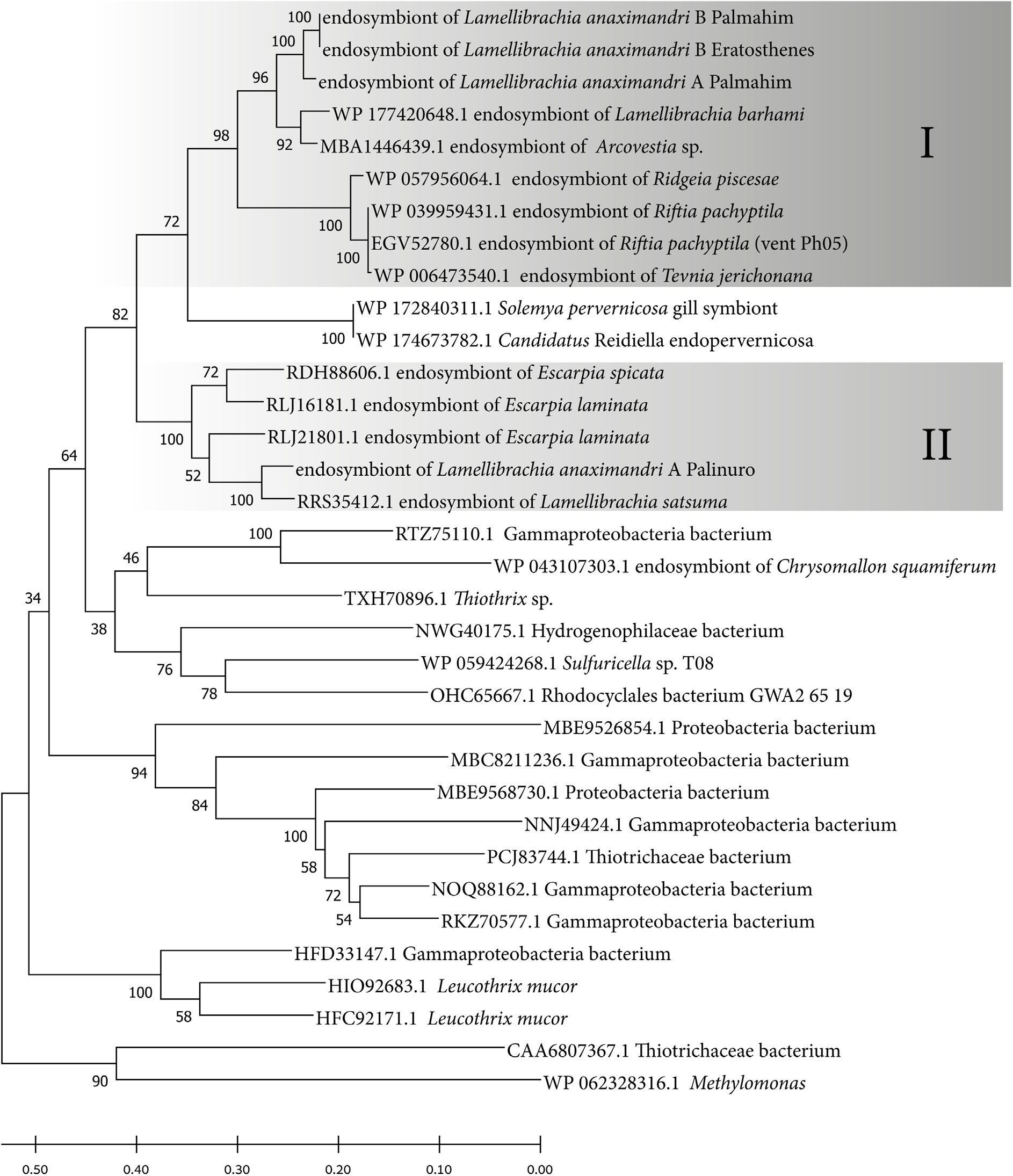
Phylogeny of the NasB (the small subunit of the assimilatory nitrate reductase). The tree was constructed using MEGAX using the LG model, and bootstrap support values are shown next to the branches. The two distinct clades of these sequences in tubeworm symbionts are accented by the grey frames.

**Figure S5:**
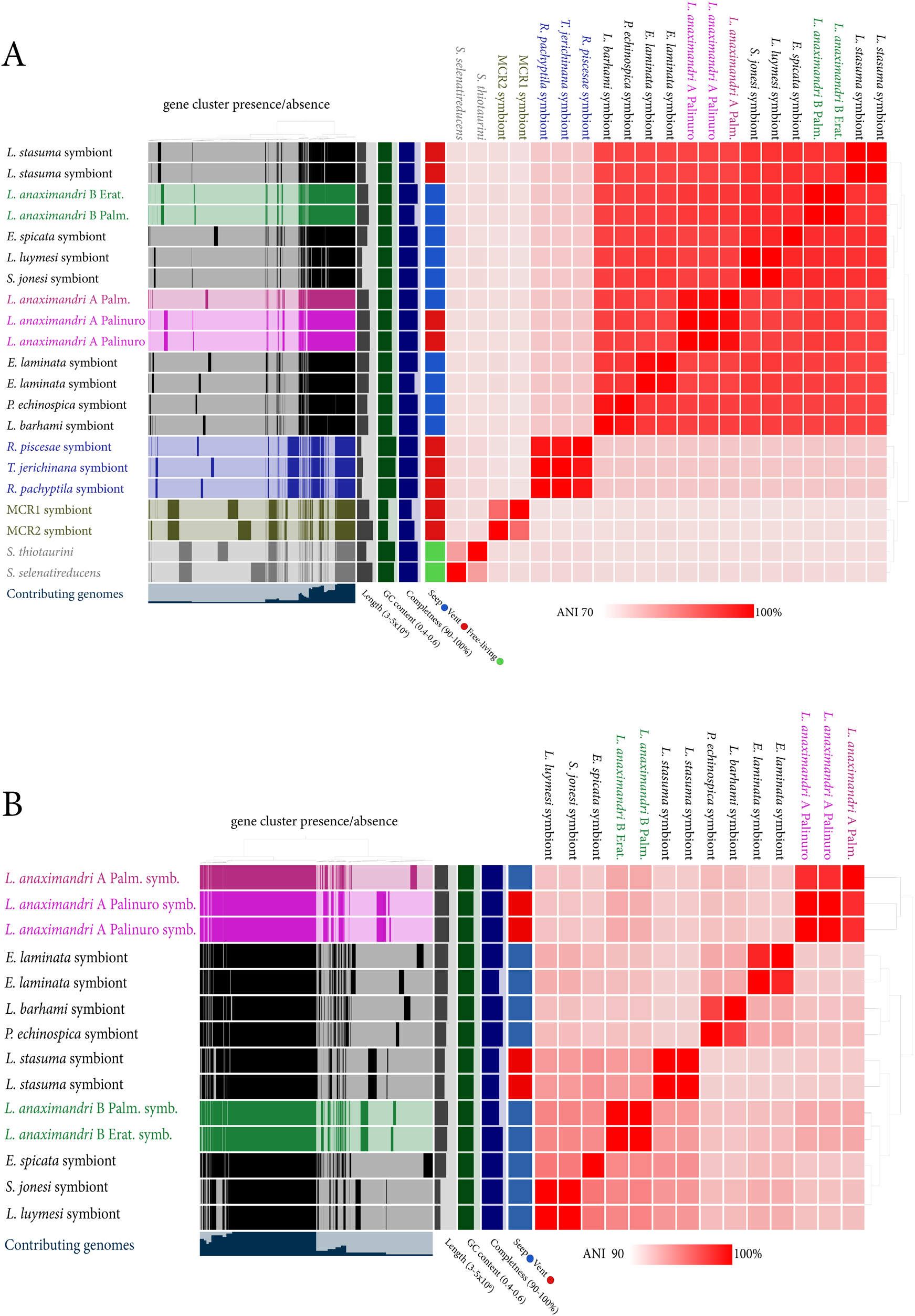
Pangenome analysis of tubeworm symbionts including the closest free-living relatives (A) and that of the seep clade only (B). Genome statistics (completeness, GC content and length), habitat metadata, average nucleotide identity heatmap and clustering are shown. Bar plots on the left represent gene cluster absence/presence and are clustered accordingly.

